# An easily adopted murine model of distal pancreatectomy for investigating immunotherapy efficacy in resectable pancreatic adenocarcinoma

**DOI:** 10.1101/2020.03.17.990903

**Authors:** Katherine E Baxter, Christiano Tanese de Souza, Lee-Hwa Tai, Pasha Yaghini, Manijeh Daneshmand, John C Bell, Brian D Lichty, Michael A Kennedy, Rebecca C Auer

## Abstract

**Background:** Although surgery provides the greatest therapeutic benefit to eligible pancreatic ductal adenocarcinoma (PDAC) patients it does not significantly improve survival for the majority of patients. Unfortunately our understanding of the therapeutic benefit of combining surgery with different treatment modalities including promising immunotherapeutics is limited by the current lack of easily adopted surgical models. The purpose of this study was to develop a surgically resectable model of PDAC in immunocompetent mice for use in preclinical investigations.

**Materials and Methods:** Surgically resectable orthotopic tumors were generated by injecting Pan02 cells into the tail of the pancreas. Fifteen days post implantation the primary tumors and tail of the pancreas were resected by laparotomy while preserving the spleen. Splenic function, tumor growth, immune phenotyping and survival were assessed following surgical resection of the primary tumor mass.

**Results:** As expected orthotopic tumor implants recapitulated many of the major histological hallmarks of PDAC including disrupted lobular structure and vascular invasion. Preservation of splenic immune cell viability and function was not associated with improved survival following surgery alone. However, pre-operative vaccination with GVAX was associated with improved survival which was not impacted by surgery.

**Conclusion:** This represents the first murine model of surgically resectable murine model of PDAC which recapitulates known pathological hallmarks of human disease in an immune competent model while allowing spleen preservation. This relatively simple and easily adopted approach provides an ideal platform to examine the efficacy of potential immunotherapy combinations for PDAC surgery patients.

## Introduction

Only 8% of patients diagnosed with pancreatic cancer today survive longer than 5 years despite aggressive interventions^1^. Although surgery can provide a clear therapeutic benefit for eligible patients, especially when combined with radiation and chemotherapy^2–4^, a sustained clinical response is rarely achieved in the majority of patients.

Our understanding of the underlying factors which contribute to disease recurrence following surgery is currently limited by the paucity of immunocompetent preclinical models of surgically resectable PDAC. The potential of such models to provide critical insights has been elegantly demonstrated in two recent publications which characterize the impact of (neo)adjuvant chemotherapy treatment on the immune system and tumor growth following surgical resection^5,6^. However, widespread implementation of this model may be limited by the challenges associated with acquiring and maintaining the transgenic mouse strain and the in vivo electroporation procedures needed for initiating tumor growth.

In this report we describe the development of a surgically resectable orthotopic model of PDAC which reflects many of the hallmarks of human PDAC pathology which only requires the implantation of the well characterized, widely available Pan02 cell line. Additionally, we demonstrate that this model can be used to investigate the effects of surgery on a clinically relevant immunotherapy.

## Materials and Methods

### Cell lines

Pan02 murine pancreatic adenocarcinoma cell line^7^ was obtained from Drs Carolina Ilkow and John Bell (OHRI, Ottawa), and was maintained in DMEM (Corning Cellgro 10-013-CV) supplemented with 10% FBS (Seradigm #1500-500). B78H1GM cells were obtained from Dr Elizabeth Jaffee (Johns Hopkins, Baltimore). Cells used during this study were less than passage 30 and confirmed to be free of mycoplasma contamination by Hoescht staining (Thermofisher Cat#62249) as per manufacturer’s protocol.

### Animals

Female C57BL/6 mice 6 weeks of age were purchased from Charles River Laboratories. Male animals were not included in the current study as sex is not anticipated to impact the surgical procedure or outcomes. Animals were housed in pathogen-free conditions and all studies performed were in accordance with institutional guidelines at the Animal Care Veterinary Service facility of the University of Ottawa (Ontario). The Canadian Council on Animal Care and the Animal Care Committee of the University of Ottawa approved this study. Mice were euthanized by intraperitoneal (IP) injection of Pentobarbital Sodium (65mg/kg).

### Tumor implantation

Routine perioperative care (including anesthesia and pain management) was performed as previously described^8^. Briefly, mice were subjected to 2.5% isofluorane (Baxter Corp.) for induction and maintenance of anesthesia. Hair was removed from the surgical site with Nair™ or by shaving. For all surgical procedures, the surgical site and surrounding areas are prepared with Betadine® surgical scrub, 70% alcohol, and Betadine® surgical paint. Following a mid-line incision (1-3 cm) the tail of the pancreas was identified and isolated using a cotton swab moistened with saline. A 30G ½ inch needle (Hamilton, Reno 7655-07, Model 1705RN syringe, Hamilton 7803-07, length 0.625” needle) was used to inject Pan02 cells in a total volume of 10 μl of PBS into the tail of the pancreas. Following removal of the needle the injection site was inspected to ensure no leakage prior to cleaning the injection site with saline and returning the pancreas to the abdominal cavity. The underlying muscle and skin are then sutured using 5.0 Polysorb™ absorbable suture (Covidien) or 9mm stainless steel wound clips (BrainTree Scientific) respectively. The clips were removed 7 days following tumor injection. <u>(Supplementary Video 1)</u>

### Surgical resection

Primary tumors were resected by laparotomy on day 15 (1×10^5^ cells implanted) or day 20 (1×10^3^ and 1×10^4^ implanted cells). Briefly, mice are anesthetized and prepared as described above. A 1-1.5 cm midline incision is then made starting at the umbilicus and cutting upward towards the sternum through the skin and muscle along the linea alba to ensure minimal bleeding. The tail of the pancreas and adjacent area is then located and isolated with sterile saline soaked cotton swab. The tumor along the splenic portion of the pancreas is isolated from the splenic artery using sutures and removed, keeping the mesenteric pancreas intact using sutures (5.0 Polysorb™ absorbable sutures (Covidien) or PDS II suture (Ethicon Z423)). Any bleeding observed in the abdomen was located and sutured (arteries) or cauterized to limit mortality post-surgery. The surgical incision was closed as described above using Polysorb™ sutures and staples. All mice undergoing surgery received pre- and post-operative care including daily wellness and injections of buprenorphine (0.05mg/kg) subcutaneously (SC) 1hr pre-op and every 8 hours for 2 days following surgery. <u>(Supplementary Video 2)</u>

### Vaccination

As per Leao et al^9^, regulatory T cells were depleted one day prior to vaccination and was achieved with a single dose of 50μg anti-CD25 (Clone PC61.5, Biolegend) with low dose cyclophosphamide (100mg/kg, The Ottawa Hospital, Ottawa, ON) delivered in a 200μl volume *i.p.* GVAX, consisting of 2e6 B78H1GM and 2e6 Pan02 (irradiated at 60Gy) combined 1:1, was injected subcutaneously into both hind limbs at day 13 following tumour implantation.

### Histology and Immunohistochemistry of tumor samples

Pancreatic tumors were collected following surgical resection and at endpoint. Collected tissues were fixed in formalin (Fisher Chemical, SF100-20) for 48 hrs prior to paraffin embedding and sectioning by the University of Ottawa Histology Core Facility. Sample sections were stained by hematoxylin and eosin (H&E) for histological analysis or were stained by immunohistochemistry for CD3 (Rabbit Anti-CD3,Abcam ab5690), CD4 (Rabbit Anti-CD4, Abcam ab183685) using the Dako EnVision+ System (K4000), CD8 (eBioscience 14-0808-80) using biotinylated Rabbit Anti-Rat IgG (ab6733). Both systems were visualized with ImmPACT™ DAB (SK-4105) and counterstained with hematoxylin. All samples were analyzed by a certified pathologist for tissue identification, analysis and validation.

### Flow cytometry of tumor samples

Pancreatic tumors were collected following surgical resection, and were placed in dissociation media consisting of RPMI supplemented with 10% FBS, Collagenase I and DNase IV. Following manual dissociation and incubation for 1 hr, the resulting mixture was passed through a 70μm filter. The tumor cells were subsequently stained with CD45-BV786 (BD Biosciences 564225), CD3-AF700 (BD Biosciences 557984), CD8α-PE-CF594(BD Biosciences 562283), CD4-PE (BD Biosciences 553730), CD25-FITC(BioLegend 102006) or CD25-PE(eBiosciences 12-0251-81), Foxp3-APC(eBiosciences 7-5773-80), CD11b-APC(eBiosciences 17-0112-81), CD11c-PE(eBiosciences 12-0114-81), CD69-FITC(eBiosciences 11-0691-81), CD49b-APC(eBiosciences 17-5971-81), CD122-PE(eBiosciences 12-1222-81), PD1-APC(BD Biosciences 562571), CD19-PE(BD Biosciences 557399), Gr-1-FITC (eBiosciences 11-5931-82).

### Peritoneal weight

The entire contents of the peritoneal cavity including tumor tissue was removed and weighed and normalized to body weight (peritoneal weight / total body mass x 100%) to determine the overall tumor burden present.

### Splenocyte viability and cytokine production

The viability and functionality of splenocytes isolated 20 days following partial pancreatectomy were assessed in mice receiving an initial injection of 1×10^4^ Pan02 cells. Briefly, viability of splenocytes isolated as previously described^10^ was determined by enumerating the proportion of Trypan-blue (Sigma T8154) excluding cells as a percentage of the total number of splenocytes. IFNγ and TNFα production were assessed by flow cytometry following stimulation of 1×10^6^ whole splenocytes with 0.1μg/ml of phorbol 12-myristate 13-acetate (PMA, Sigma P8139) and 1μg/ml of Ionomycin (Sigma I0634) for 4 hrs in the presence of 1μg/ml GolgiPlug™ (BD Pharmingen, 555029) as previously described^10^. Following incubation cells were incubated with fluorochrome labelled antibodies, and then were permeabilized and fixed with Cytofix/Cytoperm (BD Pharmingen, 555028) and stained for intracellular cytokines. Antibodies used include: CD3-PerCP (Biolegend 100326, clone 145-2C11), CD8α-FITC (BD Biosciences 553031, clone 53-6.7), IFNγ-APC (eBiosciences 17-7311-81, clone XMG1.2) and TNFα-PeCy7 (Biolegend 506324, clone MP6-XT22).

### Statistical Analysis

One-way analysis of variance (ANOVA) with Tukey post hoc tests or Student’s T test were performed for all data. A P-value of <0.05 was considered significant. Survival was plotted as a Kaplan-Meier survival curve and statistical differences assessed by log-rank tests.

## Results

### Orthotopic injection of Pan02 cells into the tail of the pancreas results in localized pancreatic tumor with development of metastatic peritoneal disease

To develop a surgically resectable murine model of PDAC which recapitulates human disease we first sought to determine the optimal number of Pan02 that could be orthotopically implanted into the pancreas and yield a resectable primary tumor. Total cell numbers of 1×10^3^, 1×10^4^ or 1×10^5^ were injected into the tail of the surgically exposed pancreas and the tumor burden examined at 15 and 20 days post seeding (Figure 1 and Table 1). Notably, tumour outgrowth was present at the injection site in the majority of mice at all injection doses (Figure 1B-D). Peritoneal disease was also visibly absent in the majority of mice receiving 1×10^3^ or 1×10^4^ cells by day 15, a finding which most closely reflects the disease status of patients eligible for surgical resection. In contrast, the highest dose of 1×10^5^ had a consistently higher tumor burden with 28% of mice (5/18) having disseminated peritoneal disease as early as day 15 (Table 1). In order to determine if leakage of malignant cells from the primary injection site was contributing to the development of the observed peritoneal disease, we performed a resection of the pancreatic tail immediately following tumor injection. Pancreatic tumor outgrowth was not observed in either the pancreatic remnant or peritoneal cavity suggesting peritoneal disease was not a result of increased cell leakage under these conditions (Table 1 “Immediate Resection”).

**Figure 1:**
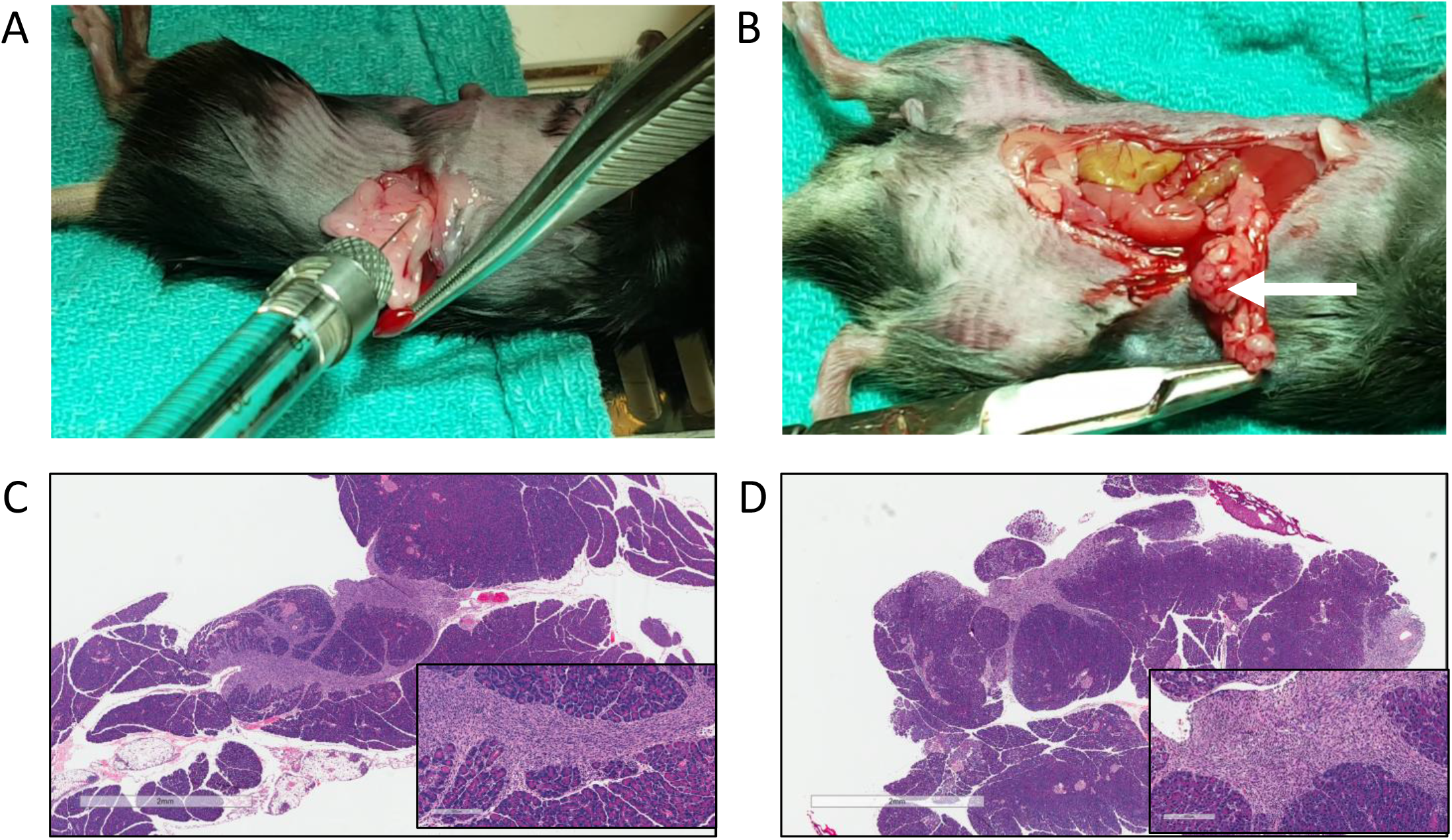
Injection of Pan02 cells into the tail of the pancreas. (A) A single cell suspension of Pan02 cells (10μl) were injected into the tail of the pancreas using a Hamilton syringe. (B) Representative images of pancreatic tumor burden (white arrow) at time of surgical resection. The tail of the pancreas was excised as described in the methods, fixed in paraffin and stained with H&E to reveal tumor burden after injection of (C) 1×10^4^, and (D)1×10^5^ Pan02 cells (30X magnification scale bar = 2mm, inset 200X magnification, scale bar = 200μM).

**Table 1:** Incidence of primary and metastatic disease in the surgical model

|  |  | **Immediate**<br>**Resection** | **$1 \times 10^3$** | **$1 \times 10^4$** | **$1 \times 10^5$** |
| --- | --- | --- | --- | --- | --- |
| **At time of**<br>**surgery** | Orthotopic Tumor | 0/5 | 4/5 | 11/12 | 18/18 |
|  | Peritoneal disease | 0/5 | 0/5 | 1/12 | 5/18 |
| **Post-surgery** | MRD outgrowth | 0/5 | 2/5 | 4/5 | 5/5 |
|  | Peritoneal disease | 0/5 | 2/5 | 4/5 | 5/5 |

### Hallmarks of PDAC are evident in the orthotopic Pan02 tumor model

Next, we examined whether the primary tumor outgrowth was reflective of human disease by histological examination of resected tumors. A trained pathologist identified 5 of the 9 established histological features associated with invasive carcinoma as described by Hruban and Fukushima^11^ in 9 of 10 animals suggesting that orthotopic injection of Pan02 cells into the tail of the pancreas results in a pancreatic disease representative of human pathology (Figure 2).

**Figure 2:**
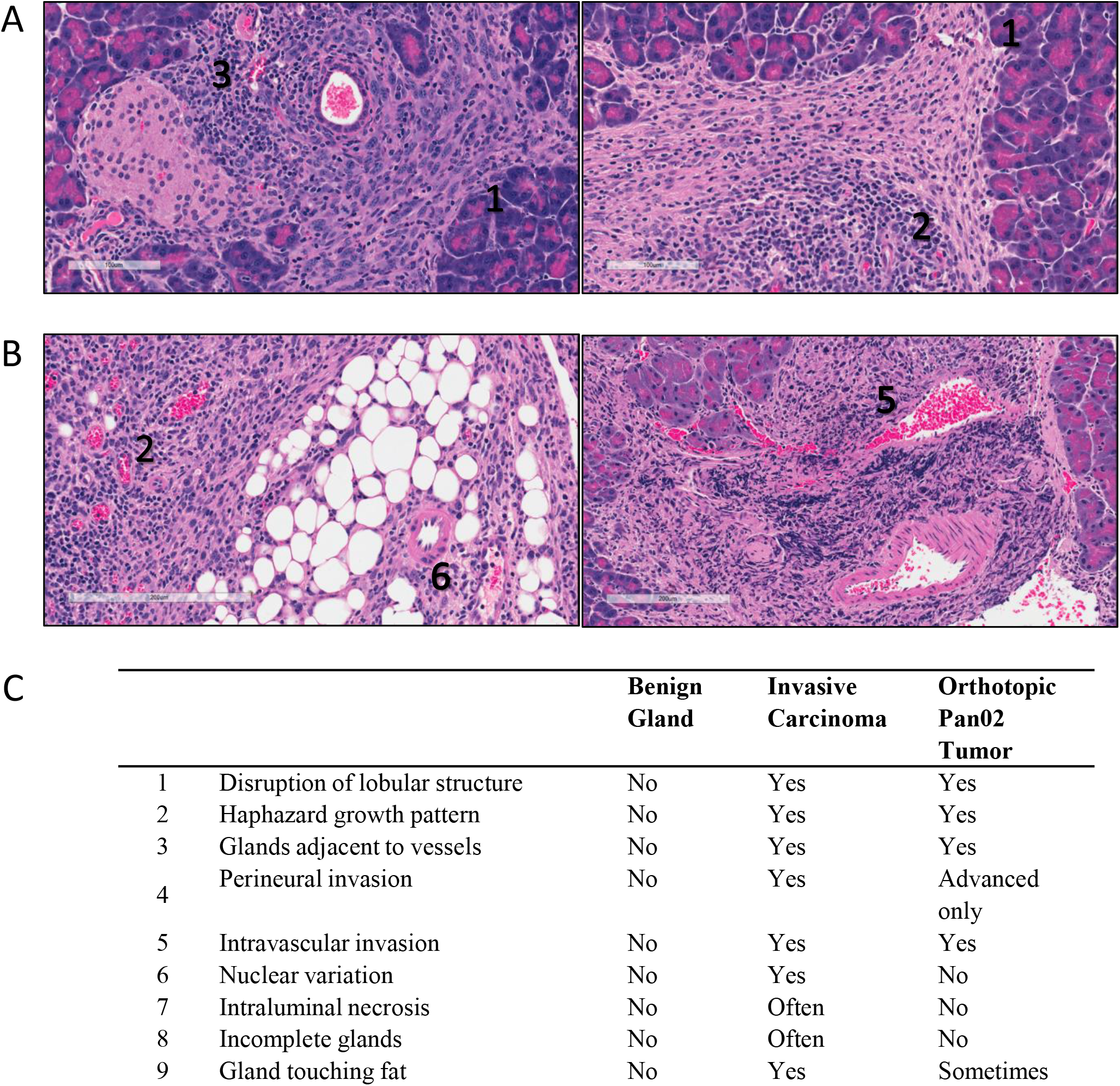
Clinical hallmarks of PDAC are evident in orthotopic Pan02 tumors. (A and B) Primary pancreatic tumors derived from Pan02 cells injected into the pancreatic tail were stained using a hematoxylin and eosin (H&E) to investigate the histopathology of the tumors. (C) Clinical hallmarks of disease compared across PDAC and the Pan02 model – adapted from Hruban et al 2007^11^. Hallmarks present in the tumors are labelled in panels A and B.

### Immune infiltrates of the Pan02 tumor microenvironment include T cell subsets

Lymphocytic infiltration is strongly correlated with patient outcomes in PDAC^12–14^ therefore we also sought to characterize the immune cell populations present within the orthotopic tumor (Figure 3). Interestingly, CD3^+^ cells were consistently observed within the cancerous regions of the pancreas but were rarely found within the healthy pancreatic tissue (Figure 3A, compare tumor (T) with healthy pancreatic tissue (P)). Subsequent staining revealed that both CD4^+^ and CD8^+^ cells contribute to the CD3^+^ population found within the TME. Flow cytometric analysis of dissociated tumors further revealed that CD4^+^ and CD8^+^ T cells were present in nearly equal proportions suggesting the presence of an active immune response to the growing tumor (Figure 3B). Interestingly, a significant proportion of the infiltrating CD4+ population also expressed markers of regulatory T cells suggesting the presence of an immunosuppressive tumor microenvironment (TME) (Tregs, CD4^+^ Foxp3^+^ CD25^+^, p=0.0004 over control mice, Figure 3C). Consistent with this observation CD8^+^ T cells found within the TME consistently exhibited increased expression of PD-1 compared to circulating T cells (p=0.01) (Figure 3D). Further examination of the immune infiltrate also revealed the presence of additional cell populations with potentially suppressive activity including myeloid derived suppressor cells (CD11b^+^ Gr1^+^) and a large number of CD19^+^ B cells which could possibly represent regulatory B cells^15^ (Figure 3B).

**Figure 3:**
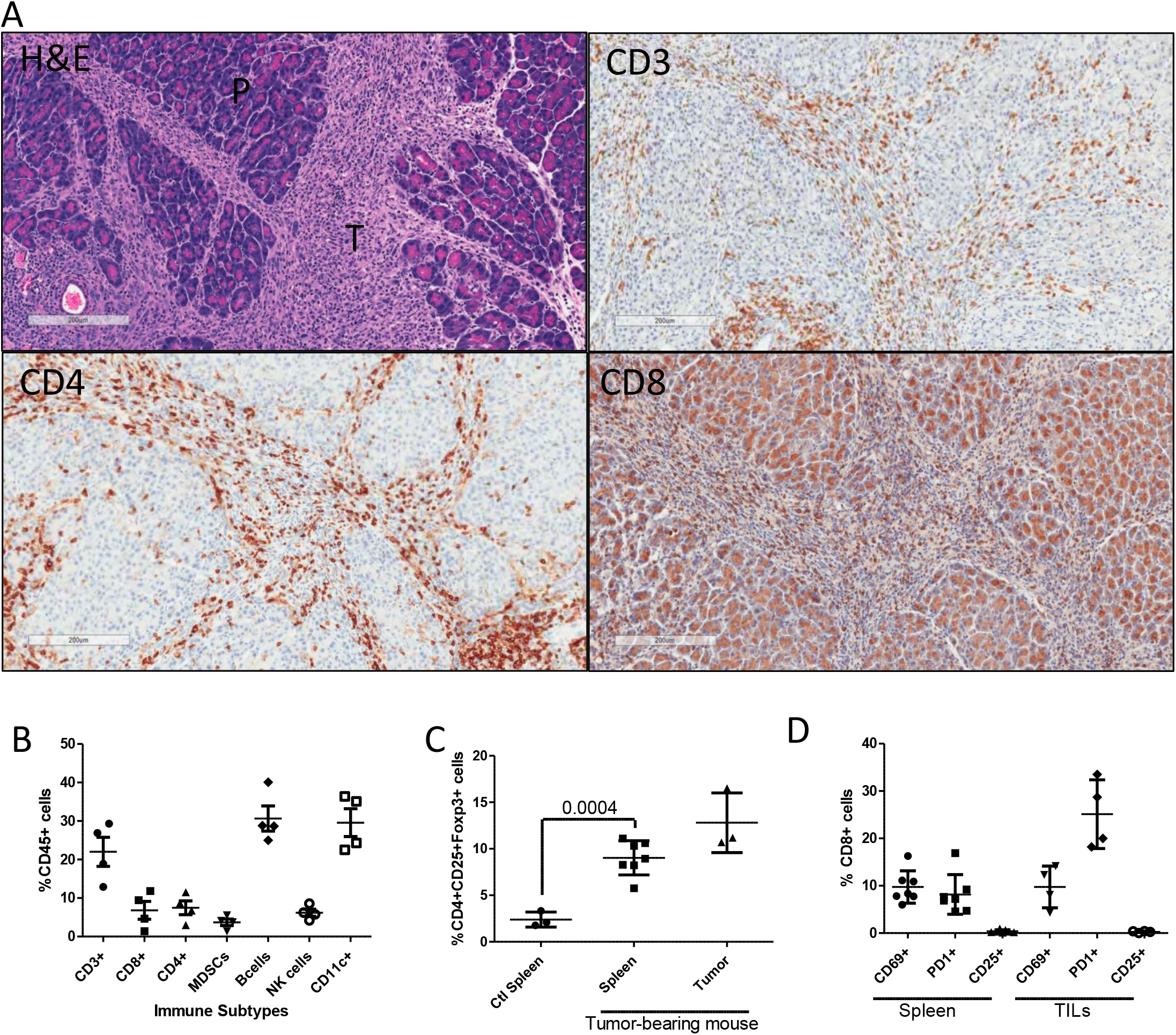
Immune cells infiltrate primary Pan02 tumors. Tumor IHC and flow cytometry was performed as described. (A) Serial sections of tumor (T) and surrounding healthy pancreas (P) collected 20 days post orthotopic injection stained by H&E, anti-CD3, anti-CD4, and anti-CD8 as marked. Images representative of all mice (n=5). Magnification 200X, scale bar = 200 μM. (B) Flow cytometry analysis of CD45^+^ tumor infiltrates at day 20 (n=4). (C) Regulatory T cell infiltrate into tumors and circulating in tumor-bearing mice as compared to a healthy control (n=3-7). (D) Activation of CD3^+^ CD8^+^ cells within the spleen (n=7) and tumors(n=4) of tumor bearing mice at day 20 post injection.

### Peritoneal disease and metastatic spread are evident following surgical resection

Having characterized the conditions necessary for establishing resectable orthotopic tumors we next aimed to determine whether this model could be used to identify parameters contributing to the success of surgical intervention. Surgical resection of the primary tumor on day 15 or 20 post injection for mice receiving either 1×10^5^ or 1×10^4^ Pan02 cells respectively ensured the resection could be performed in the absence of visible peritoneal disease, splenic metastases, or extensive adherence to organs as is generally required in the clinical setting^16^(Figure 4A and Table 1). However, despite surgical resection of the primary tumor, disease was evident in both the pancreatic remnant and disseminated throughout the peritoneal cavity by day 40 (Figure 4B and Table 1). The total disease burden, as quantified by normalizing peritoneal weight to total body mass, was significantly increased compared to a control cohort (Figure 4C, n=5, 21% body weight vs 23.1% body weight, p= 0.04). Interestingly, 50 % (4/8) of mice developed ulcerating lesions within 1 week of surgical resection of the primary tumor. These lesions were characterized by hair loss, skin erosion and full thickness abdominal wall defects. Further investigation identified leakage from the pancreatic remnant when Polysorb™ absorbable sutures were used to tie off the proximal pancreatic duct. This led to pancreatic fistulisation through the abdominal wall (Figure 4D). The use of PDS II sutures, which are commonly used in partial pancreatectomies^17^ completely prevented the occurrence of pancreatic fistulas in subsequent surgical cohorts (0 /13 animals) (Table 2).

**Figure 4:**
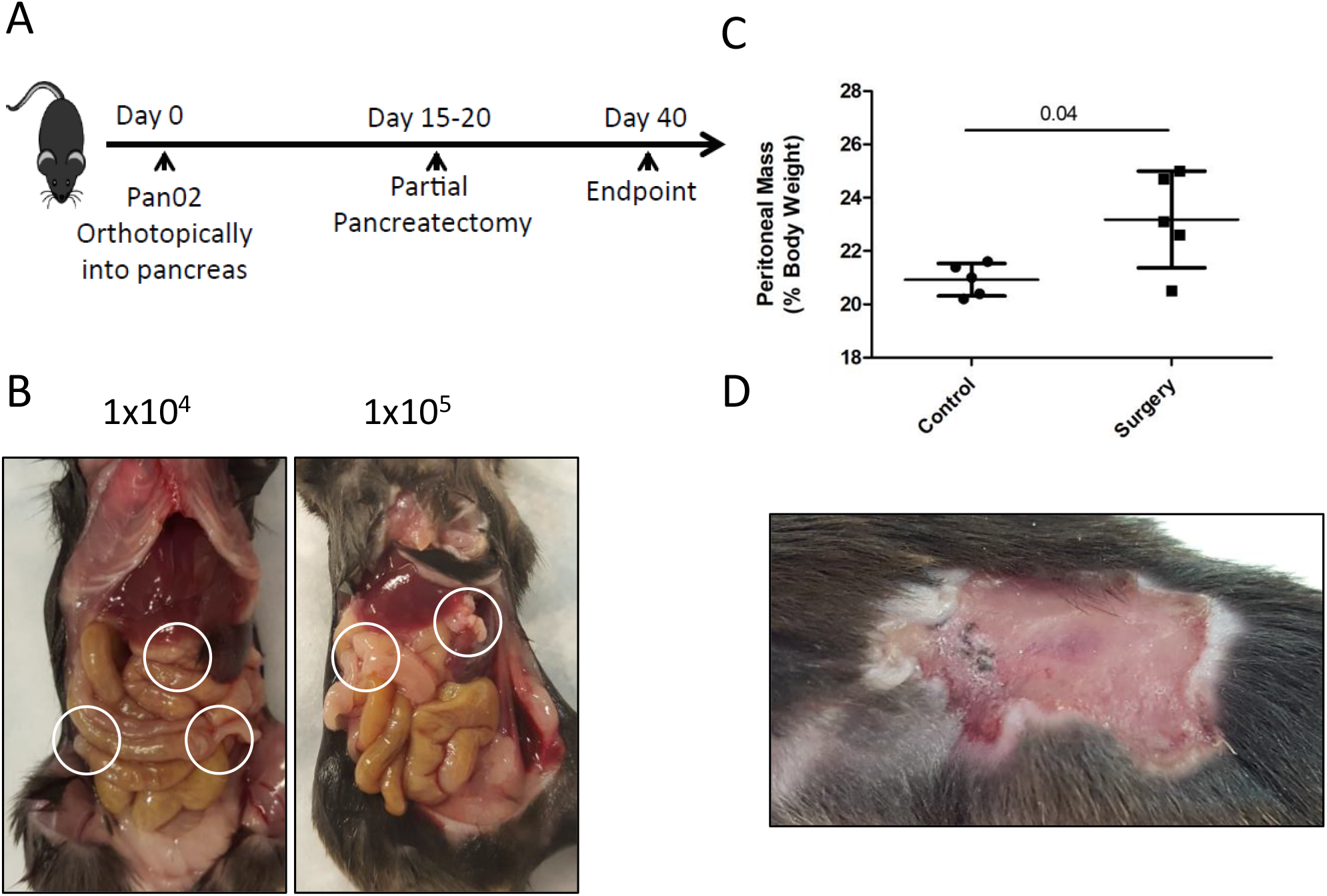
Orthotopic model is amenable to tumor surgical resection. (A) Timeline for partial pancreatectomy to resect the primary pancreatic tumor. (B) Peritoneal metastases are visible 20 days post-surgery at both initial Pan02 cell concentrations (n=10 per group). Representative images of the peritoneal cavity at 20 days post resection. (C) Weight of the peritoneal cavity normalized to the total mass of the mouse 20 days post-surgery (n=10 per group). (D) Representative image of lesion occurrence post surgery using absorbable sutures.

**Table 2:** Occurrence of skin lesions following partial pancreatectomies

| **Suture** | **Occurrence of lesions** |
| --- | --- |
| 5.0 absorbable suture | 4/8 |
| PDS II suture (Ethicon Z423) | 0/13 |

### Spleen viability and cytokine secretion are intact following surgery

In addition, we also sought to examine whether splenic immune function, in particular viability and T cell functionality, could be maintained following surgery in this model since clinical evidence supports a beneficial role of spleen preservation in improved outcomes and decreased complications^18–21^. Histological examination of the spleen indicates that surgical resection of the primary tumor does not noticeably impact splenic architecture as the white and red pulp, marginal zones and follicles were still present (Figure 5A, n=10 total number of mice examined). However, an increased splenic weight and an accumulation of erythrocytes within the red pulp were observed in mice following resection (Figure 5A and B). The apparent accumulation of blood is consistent with a reduced drainage through the splenic vein which was severed as part of our tumor resection procedure. However, despite evidence of blood accumulation, there was no significant difference in splenocyte viability between surgery and non-surgery groups (Figure 5C, n=3 vs n=5, p=0.744). Finally, no significant difference in TNFα secretion from PMA/ionomycin stimulated T cells was evident (Figure 5D, n=3 v n=5, p=0.4) suggesting the function of CD8^+^ T cells was unaltered following surgical resection.

**Figure 5:**
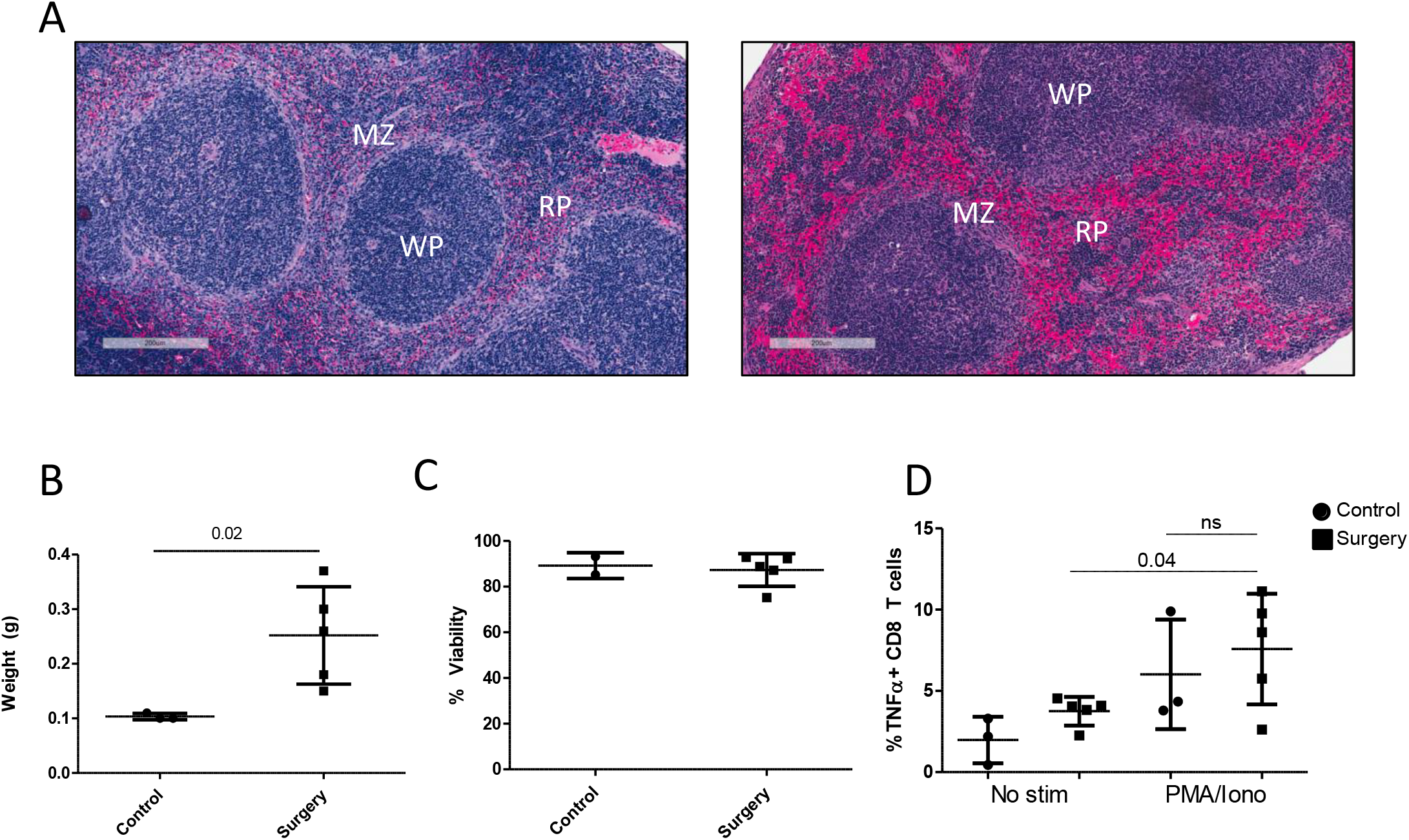
Splenic T cell function is preserved following partial pancreatectomy. (A) Histological examination of spleens from healthy mice(left), and mice 20 days following surgical resection of primary tumors(right). W=white pulp, R=red pulp, MZ = marginal zone. Scale bar = 200um. (B) Splenic weight of collected spleens. (C) Cell viability of collected splenocytes as measured by trypan blue exclusion. (D) TNFα secretion of CD8 T cells was measured by flow cytometry of whole splenocytes that received no stimulation, or PMA/Ionomycin stimulation (N=6 per group). Error bars represent standard deviation. Significance measured by Student’s t test.

### Surgical resection of primary tumor does not improve survival outcomes

Next, we sought to determine whether surgical resection could improve survival and disease recurrence in the orthotopic Pan02 model. At the time of surgery, none of the animals had visible signs of peritoneal disease and gross tumour resection was complete. Although an initial reduction in body weight occurred in the immediate perioperative period the animals undergoing surgery gained weight at a similar rate during the subsequent weeks (Figure 6A). Surgery was not found to provide any survival benefit when compared to the non-surgery group with disease evident at both the site of the primary tumor and disseminated throughout the peritoneal cavity with metastases observed in the spleen and liver (Figure 6 and Table 3).

**Figure 6:**
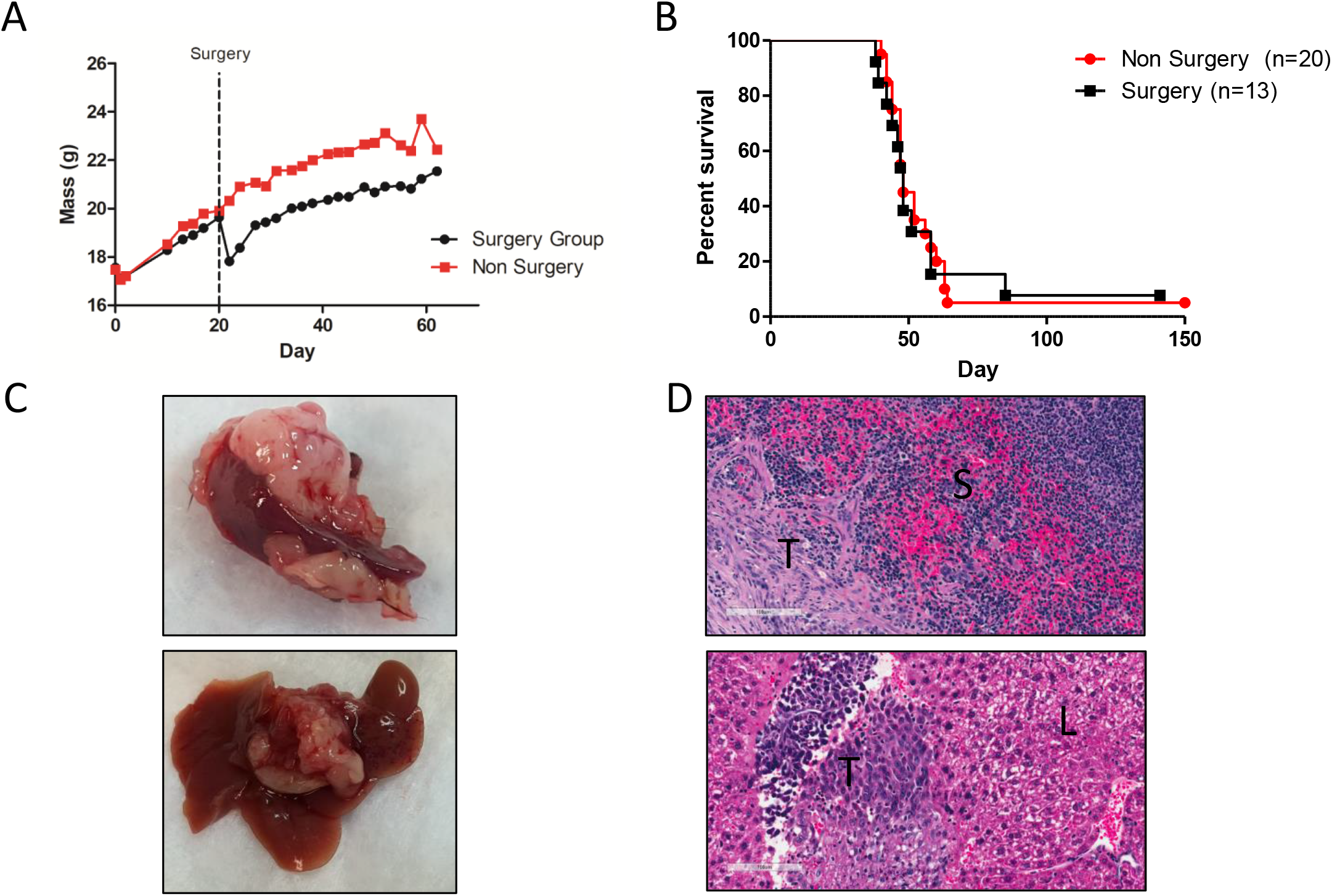
Surgical intervention has no effect on overall survival of mice. (A) Mice were weighed every day post-surgical resection of primary tumors (1×10^4^) to monitor overall health (n=7 per group). (B) Survival of mice following surgical removal of the primary tumor (1×10^5^) at day 15 (n=7 per group). (C) Tumors/metastases were specifically found to have spread to the spleen (top) and liver (bottom) of all mice at their respective endpoint (representative images). (D) The presence of tumor (T) was confirmed by H&E staining of both the Spleen (Top, S) and liver (bottom, L) (Inset magnification 200X).

**Table 3:**
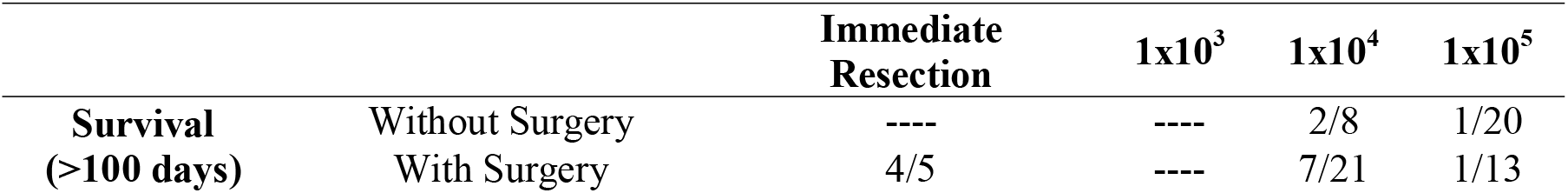
Effects of surgery on long term survival in the surgical model

|  |  | **Immediate**<br>**Resection** | **1x10<sup>3</sup>** | **1x10<sup>4</sup>** | **1x10<sup>5</sup>** |
| --- | --- | --- | --- | --- | --- |
| **Survival**<br>**(>100 days)** | Without Surgery | ---- | ---- | 2/8 | 1/20 |
|  | With Surgery | 4/5 | ---- | 7/21 | 1/13 |

### Surgery does not impact therapeutic efficacy of a clinically relevant immunotherapy

Accumulating clinical data indicates that neoadjuvant therapies can improve PDAC surgery patient outcomes^22^. However, it is unclear what impact the well-documented immunosuppressive effects of surgical stress^23,24^ may have on the efficacy of immunotherapies in PDAC. To investigate this question we examined the effects of surgery on the efficacy of GVAX, a GM-CSF producing allogeneic whole cell vaccine, currently being evaluated in PDAC surgery patients^25^. GVAX in combination with Treg depletion has previously been shown to provide a therapeutic benefit in preclinical models through immune mediated mechanisms^9,26^. To determine whether the immunosuppressive effects of surgical stress significantly impact the efficacy of GVAX we resected orthotopic tumors according to the schedule outlined (Figure 7A). In agreement with previous findings, GVAX treatment alone significantly improved survival without tumor resection (Figure 7B). Furthermore, the therapeutic benefit of GVAX was not significantly impacted by surgery.

**Figure 7:**
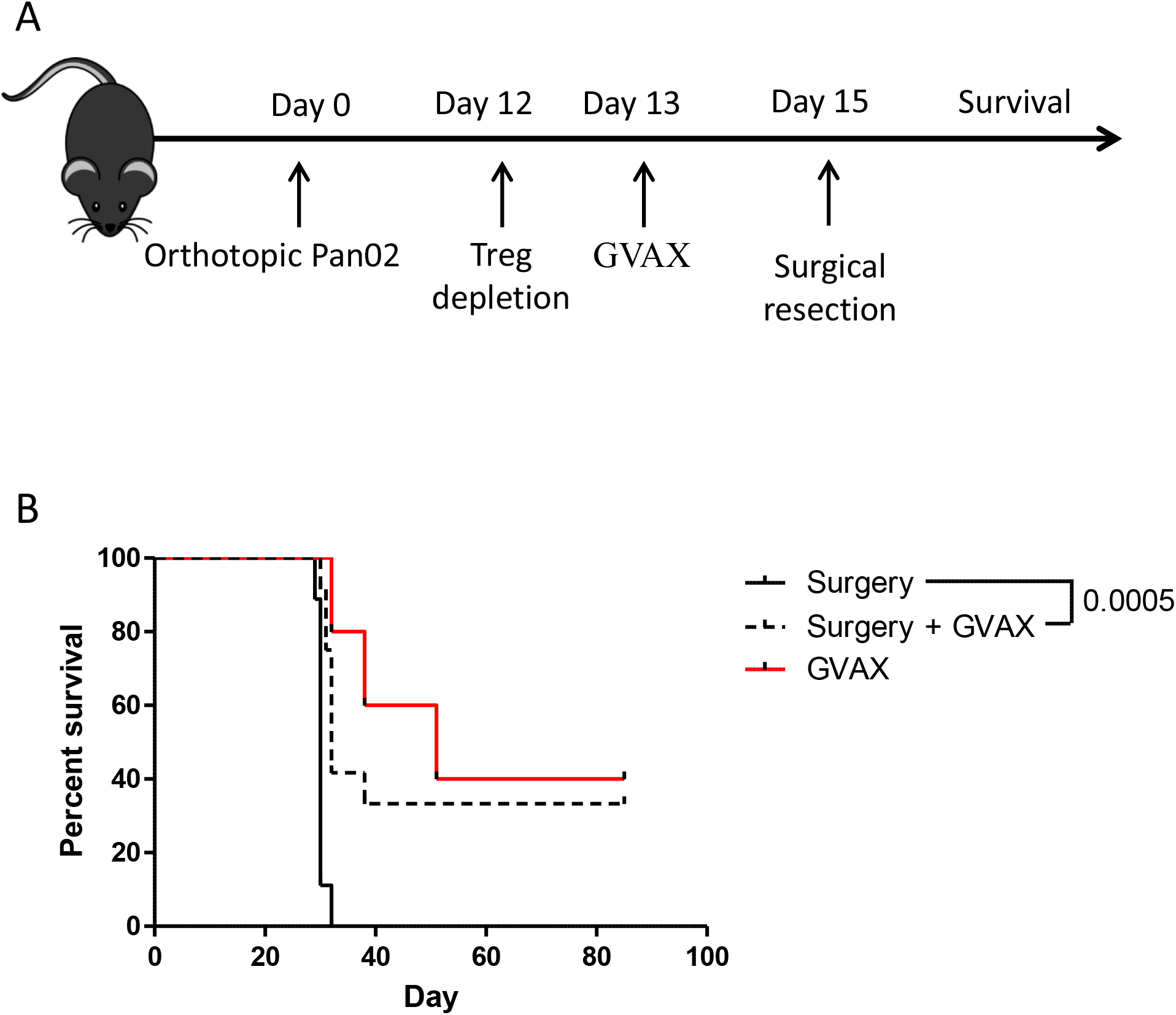
Pre-operative GVAX treatment improves survival in mice receiving a partial pancreatectomy. (A) Timeline of treatment of mice receiving 1×10^5^ Pan02 cells orthotopically into the tail of the pancreas. Total depletion of regulatory T cells was achieved with 50μg of anti-CD25 (PC61.5) and 100mg/kg of cyclophosphamide injected *i.p.* one day prior to injection of GVAX. (B) Overall survival of mice following treatment with or without surgery. N=5 in non surgery groups, and N=9 and 12 in surgery groups. Significance calculated using a log-rank test on a Kaplan-Meier survival curve.

## Discussion

Surgical resection is currently the most effective treatment strategy for patients with an operable pancreatic tumor^27^. However, despite surgical interventions PDAC patients have a very poor prognosis and it is clear that incorporation of additional therapeutic strategies is required for improving the therapeutic benefit of surgery. Unfortunately investigation of potential clinically relevant strategies in the surgical setting is limited by the lack of an easily adopted, immune competent murine model of resectable PDAC. Although Khunel and colleagues have previously developed a model that recapitulates human disease and can be successfully used to interrogate efficacy of surgery in combination with other treatment modalities^5,6^ this approach does require specialized equipment and knowledge of *in vivo* electroporation which may not be broadly available. In contrast, we sought to investigate the validity of developing an orthotopic model that requires minimal specialized knowledge beyond surgical implantation/resection techniques and can be adapted for any widely available and well characterized cell type. Consistent with previous findings^28^, we found that mice implanted with Pan02 cells develop tumors highly reminiscent of human PDAC within a relatively short time (15 to 20 days). In addition, these tumors can be resected early in disease pathogenesis, prior to visible peritoneal disease, without compromising splenic immune function. Notably, despite surgical resection of the primary tumor eventual outgrowth in both the pancreatic remnant and distant metastatic niches was evident at later time points in the majority of animals. This eventual relapse likely reflects early dissemination of tumor cells throughout the pancreatic remnant as has been previously noted in preclinical animal studies and the clinical setting.

Intriguingly, while the clinical significance of spleen preservation in PDAC is disputable, the inability of spleen preservation in surgically resected animals to provide a significant long-term survival benefit in this model suggests that other factors may limit immune mediated recognition of tumor cells necessary for disease recurrence. Indeed, immune phenotyping of the tumor microenvironment revealed an accumulation of regulatory T cells within the primary tumor and in secondary lymphoid organs. Furthermore, the increased expression of PD1 on tumor infiltrating lymphocytes may be indicative of a suppressed immune response to infiltrating tumor cells. Future studies aimed at investigating the potential role of immune subsets present at disease sites may help to clarify the role of these regulatory T cell population as well as that of the additional immune cell subsets (MDSCs and B cells) in disease recurrence. Indeed, the potential value of the current model in evaluating clinically relevant immune modulating therapies which may be combined with surgical resection was demonstrated with GVAX, a whole cell vaccine currently being explored in clinical trials^29^. Notably, we did not find a significant effect of combining neoadjuvant GVAX vaccination with surgical resection of the primary tumor despite a clear improvement in survival following treatment with GVAX alone. The similar outcomes between these groups suggests immunotherapies like GVAX may not provide additional benefits in the surgical setting despite the significant tumor debulking associated with surgical resection. It is unclear whether this is a consequence of the immunosuppressive environment associated with surgical stress^10,30,31^ or other factors associated with PDAC. However, this result provides a proof-of-concept and further emphasizes the need for models that accurately depict surgical resection of PDAC so that the timing, dose and full efficacy of combination therapies can be adequately explored prior to initiating clinical trials.

In summary, this orthotopic model recapitulates many aspects of the human disease and requires no specialized equipment or surgical procedures allowing easy adoption by other laboratories interested in assessing the effects of perioperative therapies in PDAC.

## Declarations

## Ethics approval and consent to participate

Animal studies complied with the Canadian Council on Animal Care guidelines and were approved by the University of Ottawa Animal Research Ethics Board (ME-1664).

## Consent for publication

Not applicable

## Availability of data and material

The datasets used and/or analysed during the current study are available from the corresponding author on reasonable request.

## Authors’ contributions

All authors read and approved the final manuscript. Experimental conception and design – KEB, CTS, LHT, MAK, BDL and RCA. Data Acquisition: KEB, CTS, LHT, and PY. Analysis and interpretation of data (e.g., statistical analysis, biostatistics, computational analysis): KEB, MD, MAK, and RCA. Writing, review, and revision of manuscript: KEB, MAK, and RCA. Study Supervision: L-HT, MAK, BDL, JCB and RCA.

## Acknowledgements

Pan02 cells were kindly provided by Dr. Carolina Ilkow.

